# In-Space Fabrication of Janus Base Nano-Matrix for Improved Assembly and Bioactivities

**DOI:** 10.1101/2024.03.11.584527

**Authors:** Anne Yau, Maxwell Landolina, Mari Anne Snow, Pinar Mesci, Brandon Williams, Jay Hoying, Derek Duflo, Honglu Wu, Jana Stoudemire, Rose Hernandez, Yupeng Chen

## Abstract

In-space manufacturing of nanomaterials is a promising concept while having limited successful examples. DNA-inspired Janus base nanomaterials (JBNs), used for therapeutics delivery and tissue regeneration, are fabricated via a controlled self-assembly process in water at ambient temperature, making them highly suitable for in-space manufacturing. For the first time, we designed and accomplished the production of JBNs on orbit during the Axiom-2 (Ax-2) mission demonstrating great promising and benefits of in-space manufacturing of nanomaterials.

## Content

Nanomaterial technology holds immense potential for therapeutic applications ranging from creation of biomimetic scaffolds that mimic the natural extracellular matrix (ECM) scaffolds for tissue engineering to delivery of RNAs and drugs as regenerative medicine^1,2^. Currently, many nanotechnology applications are not suitable for biomedical applications due to various issues such as complexity, and cost of nanofabrication. Scaling up these processes for commercial use can be challenging, and achieving consistent results can be difficult limiting their reproducibility. The fabrication of Janus Base Nanomaterials (JBNs), on the other hand, is simple, and the scalability and reproducibility are quick. Similar to protein crystallization^3^, the formation of JBNs on Earth is restricted due to gravity and therefore the strands formed are non-homogenous and the drug loading efficiency is less than optimal. In space, the lack of gravity can affect the sedimentation of JBNs, which can both increase homogeneity and influence their performance as drug delivery vehicles.

JBNs have emerged as a promising alternative to address the shortcomings of current therapeutic applications. These JBNs consist of small molecules mimicking DNA base pairs, self-assembled into nanotubes through hydrogen bonding and base stacking. The architecture of JBNs relies on non-covalent interactions among tens of thousands of Janus base units, each with a molecular weight below 400 Da^4,5^. These JBNs assemble via a biomimetic process at ambient temperature with minimal equipment requirements without the need for catalyst nor crosslinker during JBN formation. Due to these advantages, issues such as degradation rate, mechanical properties, and biocompatibility can be investigated easily by increasing production of JBNs on orbit.

The JBNs can further assemble with proteins to create Janus base nanomatrix (JBNms) or with RNAs to create Janus base nanopieces (JBNps)^6^. The JBNts incorporate proteins or RNAs between their bundles through positive/negative charge interactions. JBNps exhibit a rod-shaped morphology which is different from the spherical conventional delivery vehicles^7^. JBNms on the other hand are first-in-kind-injectable and multi-functional scaffolds that can be injected into tissue chips or “difficult-to-reach” locations in deep tissues. JBNm can also achieve localized drug delivery and excellent sustainability^8-10^. In Axiom Mission 2 (Ax-2), JBNms were manufactured on-orbit. The results from the Ax-2 mission helped guide the future biomanufacturing process of DNA-inspired nanomaterials, the JBN products in-space. Here, we present the results from the Ax-2 mission, launched in May 2023, to demonstrate the samples manufactured on-orbit versus lab setting, comparing JBNms with Matn1 and Matn1/TGF-β2. Utilizing current technologies, JBNs can be manufactured in a high throughput fashion, where issues such as reproducibility and scalability will not be a challenge, especially for in-space manufacturing.

JBNs are used as a starting base for the Janus base nanomatrix (JBNm) developed for cartilage repair. In space, the self-assembly of JBNs will overcome the limitations of JBN assembly process due to sedimentation^3^. Unlike many other scaffolds in the market, JBNm has room-temperature stability, quick assembly time without the need for catalysts or crosslinkers with excellent drug loading capacity^8,11^. Our first DNA Nanomaterials Therapeutics mission in the Ax-2 mission is to translate our one pot reaction to produce JBNs on Earth into operations on orbit.

We compared and observed the strands of JBNm without TGF-β2 and JBNm with TGF-β2 (**Figure 2a-b(i-iv)**) manufactured on orbit, and in lab using transmission electron microscope (TEM). We quantified the width of both JBNm strands manufactured on orbit to be larger compared to the manufactured ground samples in **Figure 2c and e**, indicating the bundling of JBNts and proteins are more efficient, which was also demonstrated in the UV-Vis spectroscopy graphs. In our characterization study, zeta potential measurement is one of the parameters that we often study to observe the formation of JBNm and determine the stability of JBNm^9^. In **Figure 2d**, we compared the zeta potential of JBNms manufactured on orbit and in lab. From previous studies, we observed that Matn1 is typically negatively charged, where it can interact with other charged proteins in the ECM through electrostatic forces and affect its interaction with calcium ions. JBNt, on the other hand, is positively charged due to its lysine side chain. The positively charged JBNt also shows that the nanoparticles are stable in physiological environment, which is an attractive property of our JBNt.

**Figure 1.**
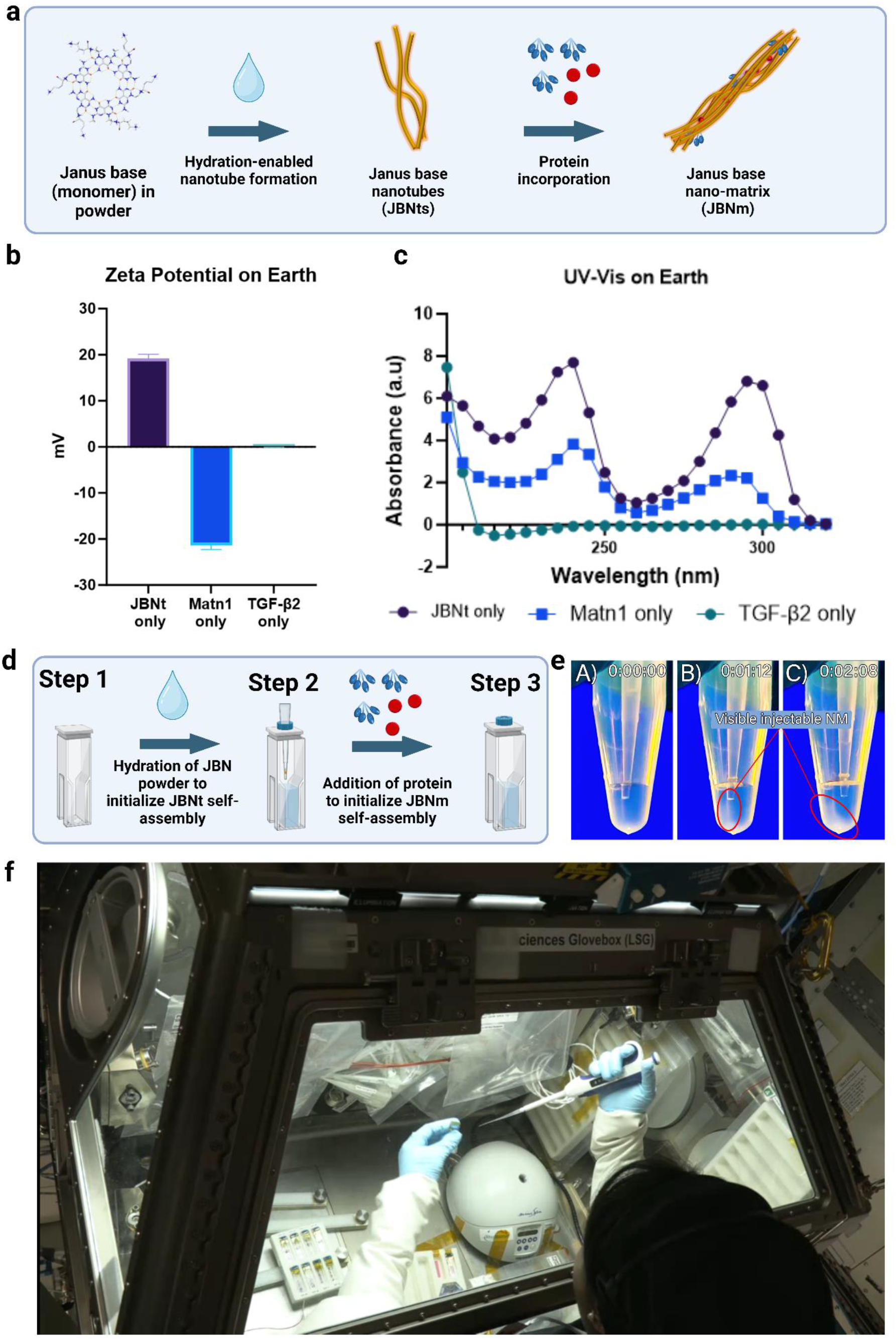
**(a)** Schematic diagram of JBNm formation with the addition of Matn1 and TGF-β2. (**b-c)** Material characterization of JBNt, Matn1 and TGF-β2 alone, UV-Vis spectrophotometer and Zeta Potential, respectively. **(d)** Schematic diagram of a one pot reaction to make JBNm done on orbit. **(e)** Screenshot clip of JBNm formation on Earth, seen to sediment on the bottom of the vial as the nano-matrix was formed one pot reaction from JBNts monomer to JBNm. **(f)** Experiments done on orbit by astronauts Rayyanah Barnawi (specialist on Ax-2 private astronaut mission) and Peggy Whitson (commander on Ax-2 private astronaut mission) in the life sciences glovebox (LSG) on the International Space Station. Image courtesy of NASA and Axiom Space.

**Figure 2.**
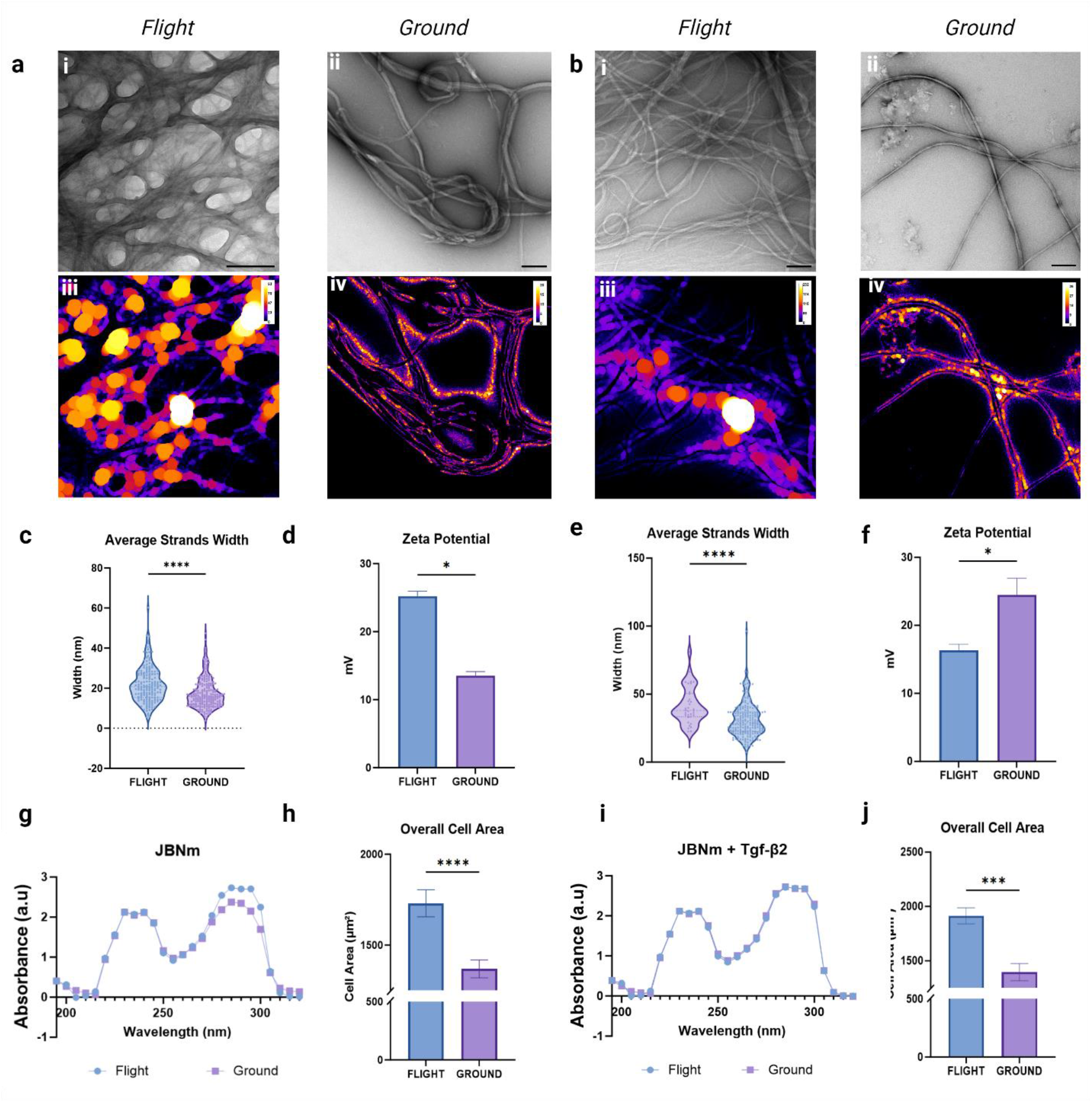
**(a-b) (i-iv)** TEM images and images processed in ImageJ of JBNm with Matn1 and JBNm with Matn1/ TGF-β2. **(c, e)** average JBNm strand width of JBNm with Matn1 and JBNm with Matn1/TGF-β2. We performed material characterization of JBNm with Matn1 and JBNm with Matn1/TGF-β2, zeta potential **(d, f)** and UV-Vis specs **(g, i)**. Lastly, we determined cartilage cell bioactivity when cultured with the samples and found that overall cell area are increased in both orbit samples **(h, j)**. Scale bar = 200 nm

Due to the increased JBNm with Matn1 properties across the board in terms of size distribution, zeta potential, the JBNm with Matn1 seem to be compatible with biological systems, which is evident in **Figure 2e**. Similarly, JBNm with Matn1/TGF-β2 also improved drug loading capacity, which is observed in TEM where JBNm bundles manufactured on orbit are larger than their counterpart, made in lab. In addition to that, the overall cell sizes in both JBNm with Matn1 and JBNm with Matn1/TGF-β2 have indeed increased when samples are cultured on cartilage cells.

The process of drug loading is crucial for the development of effective drug delivery systems^12^. The goal of drug loading is to optimize the amount of drug carried by the delivery system without compromising its stability or safety profile. We observed that the drug loading capability into JBNm, on orbit, is increased, as evident by the increase in width of JBNm + Matn1/TGF-β2 strands (**Figure 2e**), the decrease in zeta potential (**Figure 2f**), and the overall cell sizes (**Figure 2j**). While zeta potential of the JBNm with Matn1 samples made on orbit is higher than samples made in lab, the zeta potential of JBNm with Matn1 and TGF-β2 made on orbit are lower than ground samples due to the addition of TGF-β2. The zeta potential of only TGF-β2 is near neutral and the addition of JBNt to Matn1/TGF-β2 increases the surface charge of the nanoparticles in the mixture. The decrease of zeta potential in JBNm with Matn1/TGF-β2 indicated more TGF-β2 was loaded into JBNm indicating improved drug loading capacity on orbit than on Earth.

## Conclusion

JBNs are innovative DNA-inspired nanomaterials for various biomedical applications especially for growth plate fracture encouraging cartilage regeneration utilizing JBNms. Here, we analyzed in-space manufactured JBNm and Matn1 with and without TGF-β2 and found that the width of the JBNm bundles manufactured in space were larger than the JBNm bundles manufactured on Earth. We observed the JBNs had improved homogeneity and scaffold assembly in space, increasing cell bioactivity indicating low toxicity with high biocompatibility. Furthermore, tools, methodologies and SOPs developed for future missions. The mission supported by Axiom Space developed a “one-pot-reaction” methodology for nanomaterials manufacturing in space and successfully demonstrated the promise of utilizing microgravity for improved JBN assembly and bioactivities.

## Methods

### Preparation of JBNt

On Earth, the JBN self-assembly process occurs when water is added into the JBN powder. Although JBNts are instantly formed, we prepared JBNt samples on Earth by adding water into JBNt monomers. A total of two sets were prepared, one for Flight and one for Ground.

### Preparation of JBN cuvettes

First, we add water to the JBNt monomer in powder form to make a 1mg/mL solution and let it sit for 14 days at 4C. Then, we transferred 100uL of JBN (1 mg/mL) solution to 3mL cuvettes (EW-39458-60 from ColeParma). Then, we freezed and lyophilized all JBN samples (JBN S/N 1-16, 25-32) to get JBNt powdered form, except for JBNt with serial number (JBN S/N 17-24). The final weight of JBN in each cuvette should be 100 ug, except JBN S/N 17-24 would be 100uL. We then covered the cuvettes with lids (QA25 from eCuvettes). There should be a total of 32 cuvettes for JBN cuvettes.

### Preparation of Protein cuvettes

Like the preparation of JBN cuvettes, we prepared Matn1 10μg/mL proteins with and without TGF**β**2 (10μg/mL) and transferred them into cuvettes (EW-39458-60 from ColeParma) (S/N P1-P12). We then freezed and lyophilized the samples to get protein powder form and covered the top with cuvette lids (QA25 from eCuvettes). The final weight of protein in each cuvette should be 8 μg Matn1 (S/N P 1, 2, 5, 6, 9,10) and 8 μg Matn + 2 μg TGF**β**2 (S/N P 3, 4, 7, 8, 11,12). There should be a total of 12 cuvettes for protein cuvettes. ***Preparation of Water syringes***. 24 water syringes are filled with 0.5mL of DNase-free water and 12 water syringes filled with 0.9mL of DNase-free water are prepared.

### Procedures

On Day 5 of the AX-2 mission, water syringes with 0.5mL of DNase-free water were added into JBN Cuvettes S/N 1-8, 9-16 and 25-32. The solution was mixed in the cuvettes by pipetting up and down 20 times for each sample. Next, water syringes with 0.9mL of DNase-free water were added to Protein Cuvette (S/N P 9-12) and mixed with pipet and pipet tips up and down 20 times. 0.4mL of Protein solution from the protein cuvettes were then transferred to respective JBN cuvettes (S/N 29-32) and mixed up and down for 20 times. On day 10 of the AX-2 mission, water syringes with 0.9mL of DNase-free water were added into Protein Cuvette (S/N 1-8) and mixed with pipet and pipet tips up and down 20 times. Then, 0.4mL of Protein solution from the protein cuvettes were then transferred to respective JBN cuvettes (S/N 9-16, and 17-24), and mixed up and down for 20 times.

## Data availability

The datasets used and/or analyzed during the current study are available from the corresponding author on reasonable request.

## Acknowledgements

We would like to thank NIH 7R01AR072027, NIH 1R21AR079153-01A1, NSF 2025362, NSF 2234570, NASA 80JSC022CA006, DOD W81XWH2110274, Axiom Space and the University of Connecticut for funding and/or support. We would also like to thank Kevin Engelbert for his role in the government’s support of our work.

## References

1 Patra, J. K. et al. Nano based drug delivery systems: recent developments and future prospects. J Nanobiotechnology 16, 71, doi:10.1186/s12951-018-0392-8 (2018).

2 Scalia, T., Bonventre, L. Nanomaterials in Space: Technology Innovation and Economic Trends. Adv. Astronaut. Sci. Technol 3, 145–155, doi: 10.1007/s42423-020-00065-y (2020).

3 Katsuo Tsukamoto, E. F., Peter Dold, Mayumi Yamamoto, Masaru Tachibana, Kenichi Kojima, Izumi Yoshizaki, Elias Vlieg, Luis Antonio Gonzalez-Ramirez, Juan Manuel Garcia-Ruiz,. Higher growth rate of protein crystals in space than on the Earth. Journal of Crystal Growth 603, 127016 (2023).

4 Zhang, W. & Chen, Y. Molecular Engineering of DNA-inspired Janus base nanomaterials. Juniper Online J Mater Sci 5 (2019).

5 Lee, J. et al. Computation-aided Design of Rod-Shaped Janus Base Nanopieces for Improved Tissue Penetration and Therapeutics Delivery. bioRxiv, doi:10.1101/2024.01.24.577046 (2024).

6 Lee, J., Sands, I., Zhang, W., Zhou, L. & Chen, Y. DNA-inspired nanomaterials for enhanced endosomal escape. Proc Natl Acad Sci U S A 118, doi:10.1073/pnas.2104511118 (2021).

7 Griger, S., Sands, I. & Chen, Y. Comparison between Janus-Base Nanotubes and Carbon Nanotubes: A Review on Synthesis, Physicochemical Properties, and Applications. Int J Mol Sci 23, doi:10.3390/ijms23052640 (2022).

8 Zhou, L. et al. Self-assembled biomimetic Nano-Matrix for stem cell anchorage. J Biomed Mater Res A 108, 984–991, doi:10.1002/jbm.a.36875 (2020).

9 Zhou, L., Yau, A., Zhang, W. & Chen, Y. Fabrication of a Biomimetic Nano-Matrix with Janus Base Nanotubes and Fibronectin for Stem Cell Adhesion. J Vis Exp, doi:10.3791/61317 (2020).

10 Landolina, M., Yau, A. & Chen, Y. Fabrication and Characterization of Layer-by-Layer Janus Base Nano-Matrix to Promote Cartilage Regeneration. J Vis Exp, doi:10.3791/63984 (2022).

11 Yau, A., Lee, J. & Chen, Y. Nanomaterials for Protein Delivery in Anticancer Applications. Pharmaceutics 13, doi:10.3390/pharmaceutics13020155 (2021).

12 Shen, S., Wu, Y., Liu, Y. & Wu, D. High drug-loading nanomedicines: progress, current status, and prospects. Int J Nanomedicine 12, 4085–4109, doi:10.2147/IJN.S132780 (2017).

